# Random patterns of medicinal plants on a phylogeny do not imply random selections of medicinal plants

**DOI:** 10.1101/2025.02.14.638252

**Authors:** Kowiyou Yessoufou, Samaila Samaila Yaradua, Orou G. Gaoue

## Abstract

Existing studies report that medicinal plants are either phylogenetically clustered or that plant medicinal property is phylogenetically conserved. Each scenario is traditionally interpreted as evidence of non-random selection, by humans, of medicinal plants. Here, we argue that this interpretation is misleading arguing that both scenarios are simultaneously required for a correct interpretation of human selection of medicinal plants, and we propose a framework to illustrate all possible scenarios of phylogenetic patterns and human selection options.
To demonstrate the use of this framework, we conducted a survey across Katsina State in northern Nigeria, West Africa, to record plants used to treat various ailments and then employed phylogenetic comparative methods to investigate the basis for integrating a given plant in the regional pharmacopoeia.
First, we found a mixed selection pattern in organs used for medicine, perhaps mirroring the within-plants heterogeneous distribution of secondary compounds as predicted in the optimal defense theory. Second, medicinal plants used to treat most diseases follow a random distribution on the phylogeny whereas their medicinal property is convergent on the phylogeny. This pattern is plausible only when humans select preferentially less-related species for medicinal uses.
That we found a random phylogenetic structure for species used to treat most diseases would, traditionally, have been interpreted as random selections of medicinal plants. We therefore call for caution and the use of the framework we propose while interpreting phylogenetic patterns in ethnobiology.

**Societal Impact Statement:** Plants are sources of modern drugs, and traditionally used plants for medicine are the first target in bio-screening. Early discussions suggest that humans randomly select plants that they use medicinally. However, recent studies demonstrate a rather nonrandom selection pattern. Here, we argue that the conclusion of nonrandom selection is grounded on less accurate interpretation of phylogenetic patterns. We consequently propose a framework with which to interpret medicinal plant selection by human and demonstrate, using data from Nigeria in West Africa, how this framework can be used.

## INTRODUCTION

A placebo is regarded as a medicine or medical treatment carefully designed to please than to benefit the patient (Boston 1955; Frank 1975; Moreira et al. 2014; Rummun et al. 2018). Its effects are expected to act through a psychological mechanism that makes patients feel better without any scientific justification. As Beecher (1955) put it, placebo is “a psychological instrument in the therapy of certain ailments arising out of mental illness…”. Surprisingly, the magnitude of the therapeutic effectiveness of placebo is non-negligible, ranging from 30% to 60% (Beecher and Boston 1955; Frank 1975). In such context, early ethnobotanists, doubting the effectivity of traditional medicine, hypothesized that it relies heavily on placebo effects rather than on any bioactivity (see Moerman 1979; Moreira et al. 2014; Rummun et al. 2018). The implication of such position is that medicinal plants are random selections of plants from available flora.

However, plant selection for medicinal uses is increasingly demonstrated to be a non-random behavior of humans towards plants. This behavior is encapsulated in an ethnobiological hypothesis, termed non-random hypothesis (Moerman et al. 1979; Gaoue et al. 2017). The hypothesis has been variously tested, both taxonomically and phylogenetically. Taxonomically, the non-random hypothesis suggests that the number of medicinal plants in a given family is a function of the size of the family. Evidence of this taxonomic nonrandom selection is provided in several studies particularly in the Americas (e.g. in Belize, Amiguet et al. 2006; in Amazonian Ecuador, Bennett and Husby 2008; Robles Arias et al. 2020; in the US, Ford and Gaoue 2017) and in India (e.g., Kashmir, Kapur et al. 1992) with very rare attempts in Africa (e.g., Gaoue et al. 2021a; Muleba et al. 2021). Moerman et al. (1979) proposed a linear relationship between medicinal plant richness of a family and the size of the family such that larger families contain more medicinal plant species than not. Recent studies demonstrated a stronger support for an exponential function (Gaoue et al. 2021a; Muleba et al. 2021), thus raising question about previous tests of taxonomic nonrandom selection hypothesis.

One implication of the nonrandom hypothesis is that some taxa, e.g., families, are hot nodes of medicinal plants (Saslis-Lagoudakis et al. 2012), making plant families a significant predictor of plant use value (Phillips and Gentry 1993). This is because close taxa share similar evolutionary history and thus similar medicinal (biochemical) properties (Fairbrothers et al. 1975; Rønsted et al. 2008, 2012; Moerman 1991). As such, phylogeny becomes a key tool of investigation in ethnobiology and has therefore informed the formulation of a phylogeny-based nonrandom hypothesis (Saslis-Lagoudakis et al. 2011, 2012; Yessoufou et al. 2015; Gaoue et al. 2021b) which predicts that, due to shared ancestry, a sister species or phylogenetically close plant to a medicinal plant species would also be more likely medicinal than not (Yessoufou et al. 2015; Gaoue et al. 2021b).

There has been a wide spectrum of evidence supporting the power of phylogenetics comparative methods as an investigation tool not only to understand the underlying mechanisms for traditional medicine but also to advance theory-inspired and hypothesis-driven ethnobiology. For example, Yessoufou and Ambani (2021) reported that, from a phylogenetic perspective, closely related species provide similar ecosystem services, including medicinal services, whilst alien non-invasive plant species provide more services and even unique services than alien invasive species introduced to southern Africa. Also, Gaoue et al. (2021a) used phylogenetic comparative methods to demonstrate that not only medicinal plant selection is nonrandom (see also Moutouama and Gaoue 2024) but that plant organ selections are also nonrandom. Furthermore, Saslis-Lagoudakis et al. (2012) employed a phylogenetic approach to reveal how valuable traditional medicine is, in bioprospecting. They demonstrated that phylogenies can be used to provide a better understanding of the cultural patterns of plant use and guide the discovery of new medicinal plants (see also Saslis-Lagoudakis et al. 2011). Beyond local studies, a more global investigation of medicinal floras across Nepal, New Zealand and South Africa showed that medicinal plants cluster mostly within some lineages across these disparate floras, suggesting that some plant taxa are overrepresented in global medicinal floras (Saslis-Lagoudakis et al. 2012; see also Rønsted et al. 2008; Saslis-Lagoudakis et al. 2011; Weckerle et al. 2011; de Medeiros et al. 2013).

One benefit of phylogenetics based identification of nonrandom medicinal plant selection in ethnopharmacopoeia is that it makes bio-screening more focused and thus more effective than the traditional blind screening which is time-consuming and requires more financial commitments. Indeed, pharmaceutical industries, in their strides towards the identification of new molecules, tend to focus their efforts on high-throughput screening and synthetic chemistry. Such approaches could perform better if screening focused more on taxonomic groups that are more likely to have hot nodes of medicinally bioactive compound (Rønsted et al. 2008, 2012; Saslis-Lagoudakis et al. 2011, 2012).

The emerging interpretation of such phylogenetic selectivity in plant medicinal property is that human selection of medicinal plants is nonrandom (e.g., Yessoufou et al. 2015; Gaoue et al. 2021a). However, such interpretation is very limited as it fails to account for several possible scenarios. Inspired by community phylogenetics theory (see Webb et al. 2002), we propose a framework to help interpret the phylogenetic patterns of plants used in the treatment of a given disease. In this framework, we distinguish two scenarios both depending on human selection behavior (Table 1).

**Table 1.** A framework linking expected phylogenetic patterns of medicinal plants to human selection behavior and given the phylogenetic conservatism of medicinal properties.

|  | Medicinal properties are phylogenetically: |  |
| --- | --- | --- |
|  | conserved | convergent |
| <u>Human selection behavior:</u> | <u>Phylogenetic structure of plants:</u> |  |
| • **Scenario 1:** Only closely related plant species provide the medicinal properties that humans are selecting for.<br>• **Human selection behavior:** Humans filter local flora to select closely related species that provide the medicinal properties they are after | Clustered | Over-dispersed |
| • **Scenario 2:** A wide range of plant species, irrespective of their phylogenetic relatedness, provides the medicinal properties humans are after, thus giving humans the option to select the plants that provide the most competitive properties.<br>• **Human selection behavior:** Humans select the less-related plant species which provide the particular medicinal properties they are after. | Over-dispersed | Random |

In scenario 1, the plant species used to treat a given disease are more phylogenetically related than expected by chance, and as such, human selection of medicinal plants would not be random since those related plant species would preferentially be targeted for the treatment of the disease. In this scenario, if the medicinal property (or biomolecule) of the targeted plant species is shared among closely related species (that is, phylogenetically conserved), then the species used in the treatment of disease would cluster along a phylogeny (phylogenetic clustering structure; Table 1). However, if the medicinal property is not conserved, that is, convergent (implying that the property has multiple origins on the phylogeny), then the species used in the treatment of the disease would be over-dispersed (opposite of clustered) on the phylogeny (phylogenetic over-dispersed structure; Table 1). In scenario 2, the available flora offers a wide range of plant species which provide the medicinal properties that humans are selecting for, irrespective of their phylogenetic relatedness, thus giving to humans the option to select the plant species that provide, from an emic perspective (in their eyes), the most competitive medicinal properties (in scenario 1, humans have no such option but are forced to target closely related species). The concept of competitive medicinal properties here means that, by experience, humans may know which ones of the plant species that offer the medicinal property they seek, are more effective, and thus more competitive. Such group of species have competitive edge and will competitively exclude other plant species from human local ethnopharmacopoea. If such competitive medicinal property is offered by less-related species and the medicinal property is conserved on the phylogeny, then the species used in the treatment of the disease would be over-dispersed on the phylogeny. In contrast, if the medicinal property is convergent (offered by species of different origins on the phylogeny), then we would expect the species used in the treatment of the disease to be randomly distributed (Table 1) along the phylogeny (no phylogenetic structure).

New emerging diseases or pandemics are expected to occur more frequently than expected (Marani et al. 2021). In tropical regions, most regional flora are poorly explored for their medicinal potential (Soejarto et al. 2005; Gurib-Fakim 2006). Thus, testing theories or hypotheses that may accelerate success rates of biodiscovery schemes has never been this urgent. In such context, targeting woefully understudied flora in regions rich in traditional medicinal knowledge whilst conducting the phylogenetic investigation of medicinal floras, is necessary for a global establishment of the validity of the emerging hypotheses in ethnobiology (Gaoue et al. 2017; Gaoue et al. 2021b). Such investigation is required in the context where the relevance of phylogenetic comparative approach in ethnomedicine is questioned (Gertsch 2012), and phylogenetic signal in biochemistry is not always supported (e.g. Rønsted et al. 2012).

In the present study, our aim is to demonstrate how our framework (Table 1) can inform a better interpretation of the phylogenetic patterns of plant selection in ethnobiology. First, we tested the hypothesis that plant organs/products of closely related plant species are more likely to be used medicinally. Second, we tested the hypothesis that humans select preferentially plant species that are significantly closely related for medicinal uses (nonrandom selection corresponding to scenario 1 in Table 1). We tested these hypotheses using the flora of Nigeria, the most populous country in Africa with ∼ 8,000 native plant species arranged in 300 families and over 2,000 genera including 100 endemic plant species (Borokini et al. 2010; Borokini 2014). This plant diversity is used by ∼300 ethnic groups across the country (Iyiola and Adegoke Wahab 2023).

## MATERIALS AND METHODS

### Ethical considerations and consent

We received ethical approval from Katsina State Health Research Ethical Review Committee with the reference MOH/ADM/SUB/1152/1/872 (Supplemental Information – SI1). Also, we obtained written informed consent to participate in the study from all human participants (SI2).

### Study area

The study was conducted in northern Nigeria. The data was collected from six local government areas (Batagarawa, Charanchi, Mashi, Maiadua, Matazu and Dandume) in Katsina State. The majority of the people in this region demonstrated a wide knowledge of folk medicine and rely on the traditional medicine in the treatment of various ailments (Kankara et al., 2015). The people of Katsina State are mostly from the Hausa-Fulani tribe with an estimated population of 7.3 million. The study area is part of the dry Sudano-Sahelian climatic zone which is mainly dominated by scattered trees and shrubs. The annual rainfall ranges from 1000 to 1200 mm with two dry and two wet seasons (Olofin, 1985). The short dry season is dry and hot with a temperature of 40-43°C which lasts three months (March – May). This period is followed by the long raining season (May – September), the shortest dry and warm season (October – November), and a seven months-long dry season characterized by a cool Harmattan wind (November – April).

### Data collection

We conducted 11 months of field survey across six local government areas of Katsina State from February 2023 to January 2024. In each of the local governments, the district head (head of local government) was first asked to recommend 10 people who are well known within the communities as very well knowledgeable on plants and their traditional medicine. These people include traditional healers, elderly people, etc. Only people above 18 years old who are able to respond to questions independently were interviewed. A total number of 60 informants participated in the study among which 39 were men and 21 were women. The selected informants were interviewed in Hausa language at their farmlands, workplaces, homes, or markets. The interview lasted for 10 – 15 minutes, depending on the folk medicinal knowledge of the interviewee. We first explained the content and purpose of the research to the informants, and we also sought their written consent before administering the questionnaire. The ethnobotanical data of the medicinal plant was collected using a short semi-structured questionnaire. Specifically, informants were asked to list the plants they know of and tell whether the plants listed are used medicinally or not. If a plant is acknowledged as medicinal, then each informant is asked to tell the diseases the plant is used for as well as the plant organs (e.g., leaves, bark, seeds, fruits, flowers, whole plants) or plants products (e.g., gums, sap) used in the treatment of that disease. The informed consent form and questionnaire are presented in SI2.

### Data analysis

All analyses were conducted using R (R Core Team, 2021) and R scripts and data associated are available as Supplemental Information – SI3 (Yessoufou et al. 2024).

**Reconstructing phylogeny –** Using the function *phylo.maker* in the R library V.PhyloMaker (Jin and Qian 2019), we assembled the phylogeny of all plant species recorded in our survey. The Jian and Qian’s phylogenetic reconstruction uses the megatree “GBOTB. extended.tre” as backbone and this includes over 70 000 species and all families of extant vascular plants. The phylogeny reconstructed is presented as SI4 (Yessoufou et al. 2024).

**Investigating plant organ selection –** We coded organs selections as 1 (when a given plant organ or plant product is used in the treatment of a given disease) and 0 (when a given organ/product is not used). Then we tested the hypothesis that plant organs/products of closely related plant species are more likely to be used medicinally (phylogenetic signal test in plant selection). This was done using the D statistic (Fritz & Purvis, 2010). This phylogenetic signal in organ/product selection was illustrated graphically on the phylogeny using the R function contMap in the library phytools (Revell, 2012), allowing us to infer the probability of organ selection pattern in internal nodes. Data on plant organs used are presented in the Supplementary materials SI5 (see Yessoufou et al. 2024).

**Testing human selection of medicinal plants: the use of framework in table 1 –** To test the hypothesis that humans preferentially select, for medicinal uses, plant species that are significantly closely related, we employed the framework in Table 1. This framework first required that i) the phylogenetic structure of the set of species used to treat each disease be first determined, and ii) the phylogenetic signal in medicinal property be identified (Table 1). Based on informants’ medicinal knowledge, we distinguished two types of medicinal property: either a plant is ‘medicinal’ (when recognized as such by informants) or non-medicinal (when no informants recognized it as used in any disease treatment).

The phylogenetic structure was determined using NRI (Net Relatedness Index; Webb et al. 2002): NRI > 0 indicates that the set of species used to treat a given disease are phylogenetically clustered whereas NRI<0 is indicative of overdispersion and NRI=0 indicates random distribution of species along the phylogeny (Webb et al. 2002; Saslis-Lagoudakis et al. 2012). A phylogenetic clustering means that species used to treat a given disease are more closely related than expected by chance (e.g., different species belonging to the same genus or the same family are more closely related than expected by chance and as such would be clustered on a phylogeny). A phylogenetic overdispersion means that species used to treat a given disease are not closely related, that is, they belong to distantly related taxa (e.g., genera or families). A phylogenetic random pattern means a lack of phylogenetic structure implying that species used to treat a given disease are independent of phylogeny. Data required to calculate NRI is a disease x species matrix, and this is presented as SI6 (Yessoufou et al. 2024). In this data, species identified by informants as medicinally used are coded 1 and those that are not medicinal are coded 0.

Then, the phylogenetic signal in medicinal property was tested using D statistics of Fritz and Purvis (2010) as implemented in the R library CAPER (function *phylo.d*; Orme et al., 2014). D < 0 implies strong signal, D=1 means random patterns (no signal) and D > 1 means over-dispersion (medicinal property is convergent). The statistical significance of D values was tested by comparing the observed D value to 1 (random expectation). Data to calculate D are presented as SI7 (Yessoufou et al. 2024). This data includes species linked to their medicinal property (medicinal status: medicinal 1; non-medicinal 0).

Finally, we used the framework in Table 1 to determine the corresponding human behavior to the observed phylogenetic structure of plants used to treat diseases, in conjunction with the outcome of the phylogenetic signal test.

## Results

First, we tested the hypothesis that plant organs/products of closely related plant species are more likely to be used medicinally. We found support for this but only for organs such as leaves (D=0.16, P<0.001), fruits (D=0.60, P=0.003), bark (D=0.45, P<0.001) and roots (D=0.72, P=0.008; Figure 1). In contrast, there was no support for this when considering reproductive organs such as flowers (D= 0.95, P=0.376) and seed (D = 0.93, P=0.293), as well as for whole plant (D = 0.92, P=0.273), and plant products such as gum (D = 0.88, P=0.248) and sap (D = 1.75, P=0.99; Supplemental Information Figure S1; Yessoufou et al. 2024).

**Figure 1.**
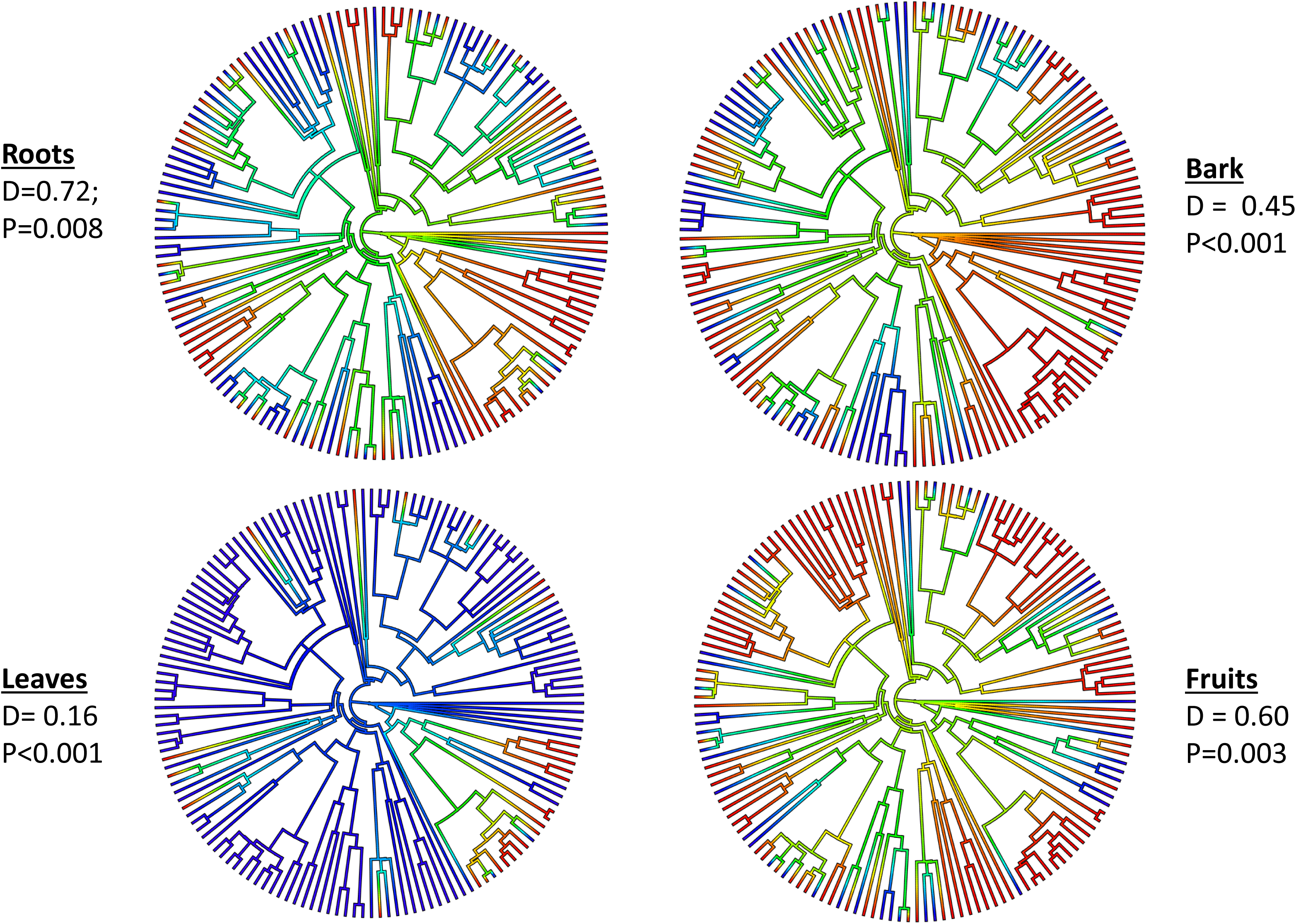
Significant clustering patterns (blue branches) of plant organ used for medicinal purposes along the phylogeny. D values are indicative of Fritz and Purvis’ (2010) phylogenetic signal in organ selection, and P values indicate significance of clustering of selected organs.

Second, we determined human selection behaviour towards plants. To this end, following the framework in table 1, we first determined the phylogenetic structure of plants used for each disease. We found a significant phylogenetic clustering pattern for only two diseases (out of 22 diseases recorded; Table S1) – stomachache (NRI=2.81, P=0.013) and gastro-intestinal disease (NRI = 2.02, P=0.03; Table S1). This finding suggests that the vast majority of diseases (91%) are treated by plants exhibiting a random phylogenetic structure (no phylogenetic structure). Then, we tested the phylogenetic signal in medicinal property. We found that medicinal property is phylogenetically convergent (D=1.047, P=0.586; Figure 2). Finally, following the framework in Table 1, an overall phylogenetic random structure of medicinal plants in conjunction with a phylogenetic convergent in medicinal property imply that humans select preferentially less-related plant species which provide the medicinal properties they are after (scenario 2 in Table 1).

**Figure 2.**
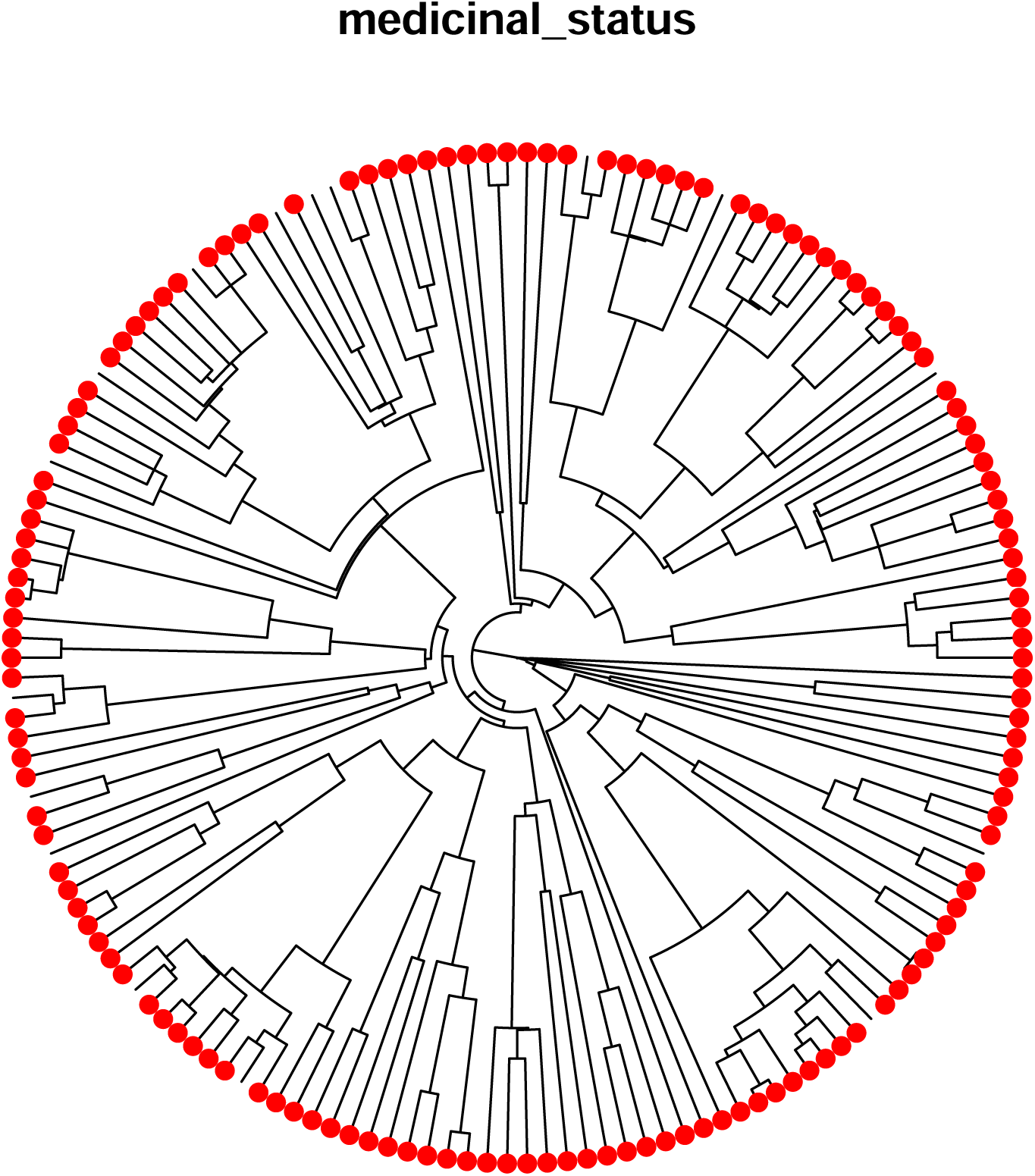
Phylogenetic convergence of medicinal status of the local flora. Red dots indicate medicinal plants. Absence of red dots indicates non-medicinal plants.

## Discussion

We found two phylogenetic patterns for plant organ selection. We found a pattern of phylogenetic signal in the use of non-reproductive organs of closely related plant species including leaves, fruits, barks, and roots. This implies that for closely related species, similar organs tend to be used medicinally. Although evidence is mounting that medicinal plant selection is phylogenetically patterned (e.g., Rønsted et al., 2012; Ernst et al., 2015; Saslis-Lagoudakis 2011,2012; Yessoufou et al. 2015; Yessoufou and Ambani 2021; Gaoue et al. 2021a,b; Moutouama and Gaoue 2024), only one study has demonstrated that even the medicinal use of plant organs is also phylogenetically patterned (Gaoue et al. 2021a). The explanation of our finding that some plant organs (those from closely related species) are used medicinally is grounded in the optimal defense theory (Strauss et al., 2004), which predicts that plant organs that are functionally important for the fitness of a given population are heavily defended chemically against attacks (e.g., herbivory). Therefore, such organs are expected to have high levels of constitutive defense (i.e., rich in secondary compounds) which would make them medicinally important for humans (Zangerl & Rutledge, 1996; Stamp, 2003; Strauss et al., 2004). Previous studies report fruits rich in toxins in several plant families including Solanaceae, Sapindaceae, and Phytolaccaceae (Cipollini & Levey, 1997). The presence of fruits in the list of organs medicinally important may *a priory* be a surprise because, if rich in secondary compounds, fruits may be less attractive or unpalatable to frugivores and dispersers. Such antagonistic interactions would compromise the dispersal and thus population fitness, unless fruits dispersal can take place via other means (e.g., dispersal by wind, water, etc.).

In contrast, we found no phylogenetic signal in the use of other organs such as flowers and seeds (see also Gaoue et al. 2021a) but also plant products such as gum and sap. This result suggests that these organs or products may be less rich in secondary compounds. The poor secondary chemistry in flower and seeds can be due to the low likelihood that these organs will be attacked as predicted in the optimal defense theory or because the plant may tolerate moderate attacks by herbivores. In their recent study, Gaoue et al. (2021a) reported a similar finding in a neighboring country to Nigeria – Benin Republic – where they showed that organs like flowers are less likely to be medicinal since these reproductive organs are mostly short-lived and therefore less available for the development of medicinal knowledge. Overall, that similar organs of closely related species are selected for medicinal uses is key for a focused bio-screening. This finding allows to target specific organs in search of new drugs or active molecules.

We further found that plant species used in the treatment of the vast majority of diseases (91%) are phylogenetically random. Does this imply that humans randomly select medicinal plants that they use? Demonstrating that medicinal plant selection is phylogenetically nonrandom is the trend in ethnobiology (e.g., Rønsted et al., 2012; Ernst et al., 2015; Saslis-Lagoudakis 2011,2012; Yessoufou et al. 2015; Yessoufou and Ambani 2021; Gaoue et al. 2021a,b; Moutouama and Gaoue 2024). These studies demonstrated that medicinal uses or medicinal properties are conserved on the phylogeny (see concept of phylogenetic conservatism in Wiens et al. 2010), which simply means that within a lineage, medicinal properties are shared among member species. Such a conservatism is traditionally interpreted as nonrandom selection, by humans, of plants they used medicinally. This implies that phylogenetic conservatism of medicinal properties predicts human behaviour towards plant selection. Here we argued we argue that this is a misinterpretation of phylogenetic conservatism of medicinal properties, and our argument is presented below.

First, there is confusion in the metrics used to test phylogenetic conservatism. Some studies used the D statistic (Fritz and Purvis 2010), a metric designed to test phylogenetic signal or conservatism in binary variables such as presence/absence of a medicinal property in a plant (Gaoue et al. 2021a,b; Yessoufou and Ambani 2021). In this approach a significant phylogenetic signal (D < 0, P<0.05) is indicative of a phylogenetic conservatism of medicinal properties. Second, other studies borrowed metrics such as NRI (Net Relatedness Index) or NTI (Net Taxon Index) from community phylogenetics (Webb et al. 2002) to infer human behaviour towards plant selection in ethnobiology. They interpreted NRI > 0 (P<0.05) as indicative of phylogenetic clustering of plants used medicinally (same for NTI>0, P<0.05; Saslis-Lagoudakis et al. 2012; Yessoufou et al. 2015; Gaoue et al. 2021b), and they interpret this as implying that humans select non-randomly medicinal plants. The misinterpretation stems from the fact that both D<0 and NRI or NTI >0 (P<0.05) are interpreted exactly the same way in ethnobiology, that is, as evidence of nonrandom human selection of medicinal plants. This is confusing since there may be phylogenetic signal in a medicinal property (D <0, P<0.05) and the group of plant species sharing that property would be phylogenetically over-dispersed (that is, NRI < 0, P<0.05; Table 1). Critically, these two metrics (D and NRI/NTI) do not necessarily tell anything about human behaviour towards plant selection, but rather tell everything about the evolutionary history of a given medicinal property (e.g., the presence of a bioactive molecule). Specifically, these metrics only reveal that a particular biomolecule or medicinal property is shared among closely related species. Therefore, demonstrating a phylogenetic conservatism (signal) in a medicinal use/property is not enough to conclude on human selection patterns: that is, the phylogenetic structure of the groups of plants used medicinally in combination with phylogenetic conservatism in medicinal property are both required before inference on human behaviour towards plants selection can be done (Table 1).

In the present study, we found that the groups of plants used to treat diseases are phylogenetically random (absence of phylogenetic pattern) whilst the medicinal property is phylogenetically convergent. Phylogenetically convergent medicinal property means that plants that are phylogenetically less related provide similar medicinal uses to humans. A phylogenetic random structure of medicinal plants in combination with a phylogenetic convergent medicinal property can only be explained by humans selecting preferentially plants species that are phylogenetically less related (scenario 2 in Table 1). Such selection can only be explained as follows. If medicinal properties are convergent, meaning that plants of multiple phylogenetic origins (i.e., plants belonging to distant lineages) provide similar medicinal uses, then humans have multiple selection options, either to select plants of the same phylogenetic origins/lineages (closely related species) or of different lineages (less related species). If for any reason, plants from different lineages are, in the eyes of peoples, more effective in the treatment of diseases, then these less related species would be preferred in the local pharmacopoeias, and as such, the groups of plants selected would exhibit a random distribution of the phylogeny. This scenario matches the pattern we observed in the present study in over 90% of the diseases recorded.

### Conclusions

Overall, we firstly found that organs of closely related plants are used for medicinal purposes. Second, we found that medicinal plants used to treat 91% of diseases are, from a phylogenetic structure perspective, random, and that the medicinal property is convergent on the phylogeny. These two scenarios imply that humans select preferentially less-related species for medicinal uses. Traditionally, the fact that we found a random phylogenetic structure for species used to treat most diseases would have been interpreted as random selection of medicinal plants. This is not correct since despite the random structure of medicinal plants on the phylogeny, humans have still selected nonrandomly plants by targeting preferentially less-related species. We call for caution while interpreting phylogenetic patterns in ethnobiology.

## Acknowledgement

We acknowledge the local communities for sharing their medicinal knowledge with us. KY acknowledged funding from the National Research Foundation—South Africa, grant number SRUG22051210107.

## Authors contribution

KY, design of the research; SSY, collected the data; KY, performed data analysis; KY, OGG performed the data interpretation; KY, OGG wrote the manuscript; KY, SSY, OGG edited the manuscript.

## Data availability

The data used for this study are available in the supplementary material of this article and also Yessoufou et al. 2024 at https://doi.org/10.5061/dryad.zw3r228gn.

## Conflict of Interest

None to declare.

## SUPPLEMENTAL INFORMATION – SI DATA

**SI 1.** Ethical approval of the study

**SI2.** Consent form and questionnaire

**SI3.** R script used for the study

**SI4.** Phylogeny assembled for the analysis

**SI5.** Plant parts used for medicinal purposes

**SI6.** Matrix used to calculate the phylogenetic structure (NRI, net relatedness index) of all plant species used in the treatment of each disease.

**SI7.** Matrix used to calculate D value for medicinal status of all species recorded.

## FIGURES

**Figure S1.** Non-significant clustering (blue branches) of plant organs and plants products used for medicinal purposes along the phylogeny. D values are indicative of Fritz and Purvis’ (2010) phylogenetic signal in organ selection, and P values indicate significance of clustering of selected organs.

## TABLES

**Table S1.** Coefficients of the net related index (NRI) of all species used in the treatments of each of the 23 diseases recorded in the present study. The highlighted rows indicate diseases treated with significantly clustered plant species. NRI=-mpd.obs.z, and P value = mpd.obs.p.

## Notes

### Competing Interest Statement

The authors have declared no competing interest.

