## Supplementary material for "Random patterns of medicinal plants on a phylogeny do not imply random selections of medicinal plants": SI1-ethical approval.pdf

Katsina State
### MINISTRY OF HEALTH
State Secretariat Complex
IBB Way Dandagoro
KATSINA


www.smoh.kts.gov.ng

GSM Lines: +23408061194975
+23408085422054
+23409092675845

MOH/ADM/SUB/1152/1/872

13th March,2024

867

**KATSINA STATE HEALTH RESEARCH ETHICAL REVIEW COMMITTEE (HREC) FULL ETHICAL CLEARANCE CERTIFICATE**

**Re: "TESTING THE SOCIAL NETWORK AND OPTIMAL DEFENSE THEORIES USING ETHNOBOTANICAL DATA FROM HAUSA ETHJNIC GROUP OF NIGERIA"**

| | | |
|---|---|---|
| Katsina HREC assigned number | - | MOH/ADM/SUB/1152/1/872 |
| Name of principal investigator | - | Dr. Sama'ila Sama'ila Yar'adua |
| Address of Principal Investigator | - | UMYU. KATSINA |
| Date of receipt of Valid Application | - | 19/02/2024 |
| Date of HREC meeting and Approval | - | 20/02/2024 |

This is to inform you that the research described in the submitted protocol, the consent forms and other participant's information materials have been reviewed and given Approval by the Katsina State Health Research Ethics Committee and accordingly by the Honorable Commissioner of Health.

Please note: this approval dates from **20/02/2024 to 20/02/2025.** No recruitment of participant into this research may be conducted outside these dates.

All informed consent forms in this study must carry the Katsina HREC assigned number and the duration of the Katsina HREC approval for the study.

If there is a delay in starting the research, please inform the Katsina HREC so that starting date can be adjusted accordingly.

No changes are permitted to the research without prior approval by the Katsina HREC except in circumstances outlined in the national code of health research ethics. http/www.nhrec.net

Katsina HREC reserves the right to conduct a compliance assessment to your research site without prior notification.

Dr Abdulrasheed Yusuf
For: Chairman, Katsina HREC
+2348067607767
