## Supplementary material for "Random patterns of medicinal plants on a phylogeny do not imply random selections of medicinal plants": SI3_R script for data analysis.R

```
rm(list=ls())
file.choose()
data_katsina <- read.table("C:\\Users\\kowiyouy\\OneDrive - University of Johannesburg\\Desktop\\UJ\\postdoc\\2023\\2024\\data\\species_list.csv",header=TRUE)
attach(data_katsina)
names(data_katsina)

#phylogeny
# load the package
library("V.PhyloMaker")
library(picante)
# input example species list
example <- read.csv("C:\\Users\\kowiyouy\\OneDrive - University of Johannesburg\\Desktop\\UJ\\postdoc\\2023\\2024\\data\\species_list.csv")
# generate the phylogeny presented in

tree.katsina <- phylo.maker(sp.list = example,
                      tree = GBOTB.extended, nodes = nodes.info.1, scenarios="S3")
write.tree(tree.katsina$scenario.3,"tree_katsina.tre3")
phylo_katsina <- read.tree("C:\\Users\\kowiyouy\\OneDrive - University of Johannesburg\\Desktop\\UJ\\postdoc\\2023\\2024\\data\\tree_katsina.tre2")
plot(phylo_katsina,cex=0.5)


######### representing uses on a phylogeny

# First read in the arguments listed at the command line
args=(commandArgs(TRUE))
library(picante)
library(ape)
library(adephylo)
library(ade4)
library(phylobase)
library(geiger)
library(phytools)
library(dplyr)

#tre_med <- read.nexus("C:\\Users\\kowiyouy\\OneDrive - University of Johannesburg\\Desktop\\UJ\\collaboration\\Gaoue\\PPP\\data\\Burned tree_March 2021.txt")
#plot(tre_med,show.tip.label=F) # read phylogenetic tree
data_med <- read.table("C:\\Users\\kowiyouy\\OneDrive - University of Johannesburg\\Desktop\\UJ\\postdoc\\2023\\2024\\data\\plant part used.txt",header=TRUE) # data that contain medicinal variables

attach(data_med)
names(data_med)

t1 <- phylo_katsina
m <- data_med
s1 <- data.frame(species=t1$tip.label)

colnames(m)
m1 <- unique(m[, c(1,10)]) # to select the corresponding column to the plant organ to represent on the phylogeny, here column 10

colnames(m1)[[1]] <- "species"

d1 <- merge(m1, s1, by="species")
head(d1)

z <- as.data.frame(d1)
rownames(z) <- z[,1]
z[,1] <- NULL

#write.csv(z, "SA/Species_with_specimens_SA.csv")
subphy <- match.phylo.data(t1, z)$phy
subdat <- match.phylo.data(t1, z)$data

names(subdat)[1] <- "x"

y <- as.numeric(subdat$x)

names(y) <- rownames(subdat)

b <- di2multi(subphy) # ensuring the tree is fully resolved

z1 <- contMap(b, y, type="fan", outline=T, fsize=0.0000000000000000000000000001,main="bark", legend=FALSE,lwd=2)

########


########### To calculate D --> phylogenetic signal test in organ selection
library(caper)

attach(data_med)
names(data_med)
rownames(data_med) <- data_med$species

phylo_katsina$node.label<-NULL
med_comp <- comparative.data(phylo_katsina, data_med, species)
d_root <- phylo.d(med_comp, binvar=Root) # to get the D value and p value for root selection
d_bark <- phylo.d(med_comp, binvar=Bark)
d_leave <- phylo.d(med_comp, binvar=Leave)
d_fruit <- phylo.d(med_comp, binvar=Fruit)
d_flower <- phylo.d(med_comp, binvar=Flower)
d_seed <- phylo.d(med_comp, binvar=Seed)
d_whole_plant <- phylo.d(med_comp, binvar=Whole_plant)
d_gum <- phylo.d(med_comp, binvar=Gum)
d_sap <- phylo.d(med_comp, binvar=Sap)


print(d_root) # D=0.7168867;P=0.008 --> not random
print(d_bark) #D =  0.4531533; P<0.001
print(d_leave) #D :  0.1624235, P<0.0001
print(d_fruit) #D :  0.601704, P=0.003
print(d_flower) # D :  0.9485527, P=0.376 ==>random
print(d_seed) #D :  0.9293483, p=0.293 ==>random
print(d_whole_plant) #D :  0.9185354, P=0.273 ==>random
print(d_gum) #D :  0.8793374, P=0.248 ==>random
print(d_sap) #D :  1.745843, P=0.991 ==>random


###########
#analysis of disease: calculate NRI
phylo_katsina <- read.tree("C:\\Users\\kowiyouy\\OneDrive - University of Johannesburg\\Desktop\\UJ\\postdoc\\2023\\2024\\data\\tree_katsina.tre")

comm <- read.table("C:\\Users\\kowiyouy\\OneDrive - University of Johannesburg\\Desktop\\UJ\\postdoc\\2023\\2024\\data\\disease_katsina.txt",header=TRUE)
comm <- as.matrix(comm)
prunedphy <- prune.sample(comm, phylo_katsina)


#######
match <- match.phylo.comm(phylo_katsina, comm) #match species on phylo with those in community
phydist <- cophenetic(match$phy)
ses.mpd.result <- ses.mpd(match$comm, phydist, null.model = "taxa.labels",
                          abundance.weighted = FALSE, runs = 999)
ses.mpd.result

write.table(ses.mpd.result,"nri_katsina2")

######## Is there phylogenetic signal in medicinal status of all plants?
data_signal <- read.table("C:\\Users\\kowiyouy\\OneDrive - University of Johannesburg\\Desktop\\UJ\\postdoc\\2023\\2024\\data\\data_signal_test_in_medicinal_status.txt",header=TRUE)
attach(data_signal)
names(data_signal)
rownames(data_signal) <- data_signal$species

phylo_katsina$node.label<-NULL
signal_comp <- comparative.data(phylo_katsina, data_signal, species)
d_med <- phylo.d(signal_comp, binvar=medicinal_status) # to get the D value and p value for root selection
print(d_med)

### representing phylogenetic signal on the phylogeny (all the plants species are used)
comm_signal <- read.table("C:\\Users\\kowiyouy\\OneDrive - University of Johannesburg\\Desktop\\UJ\\postdoc\\2023\\2024\\data\\data_comm_med_status.txt",header=TRUE)
comm_signal <- as.matrix(comm_signal)
prunedphy <- prune.sample(comm_signal, phylo_katsina)

for (i in row.names(comm_signal)) {
  plot(prunedphy, show.tip.label = FALSE,type="fan", main = i)
  tiplabels(tip = which(prunedphy$tip.label %in% names(which(comm_signal[i, ] >
                                                               0))), col=c("red"),pch = 19, cex = 1)
}
```
