## Supplementary figures and images for "Random patterns of medicinal plants on a phylogeny do not imply random selections of medicinal plants"

### Figure S1.pdf

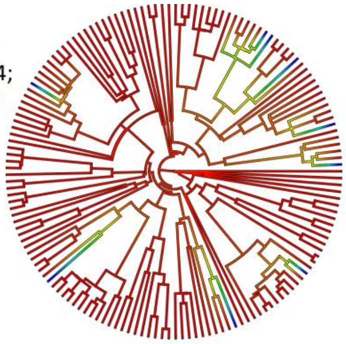

**flowers**
D =  0.94;
P=0.376

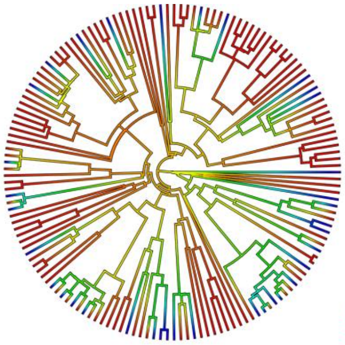

**seeds**
D = 0.93
P=0.293

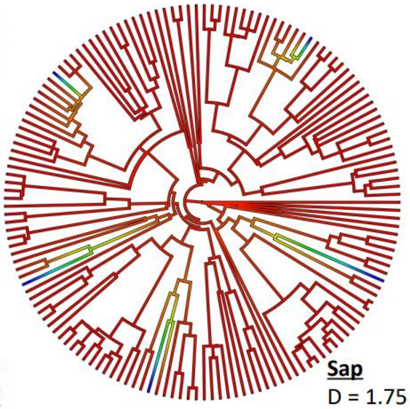

**Sap**
D = 1.75
P=0.99

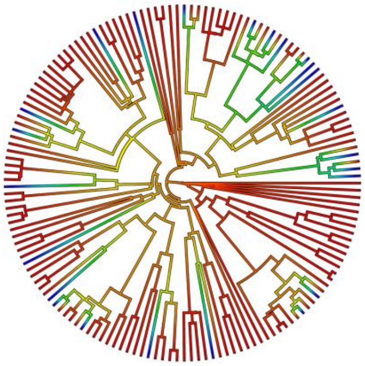

**whole plant**
D=0.92
P=0.27

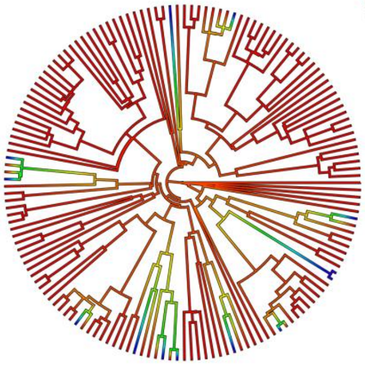

**gum**
D :  0.88,
P=0.248
